# Tagging *C. elegans* septins disrupts cytoskeletal scaffolding but not post-embryonic roles

**DOI:** 10.64898/2026.06.09.731194

**Authors:** Larry Rivenbark, Vedang Singhal, Jenna A. Perry, Amy Shaub Maddox

## Abstract

Septins are conserved polymer-forming proteins that scaffold the actomyosin cytoskeleton, its regulators, and other factors to cellular membranes. Septins also sense micron-scale curvature, bind microtubules, and establish membrane diffusion barriers. *C. elegans* is a powerful animal model to study septins’ roles because there are only two septin genes: *unc-59* and *unc-61*. In many fungal and animal cell types, septins are required for proper cytokinesis. In the *C. elegans* zygote, septins’ scaffolding roles in cytokinesis manifest during the chiral rotation of the cell cortex and the asymmetry of cytokinetic ring closure. Originally named for the uncoordinated movement exhibited by hypomorphic alleles, UNC-59 and UNC-61 are also required for normal postembryonic development, germline development, and fertility. To study *C. elegans* septins in these various contexts, we sought a fluorescent-protein tagging strategy that minimally perturbed septin function. We examined strains in which GFP, mKate2 or wrmScarlet had been inserted at the *unc-59* locus, or coupled to *unc-61b/c* at an exogenous locus, to encode fluorescently tagged fusion proteins. We compared these tagged septins to classical hypomorphic alleles, and to new null alleles. Null alleles phenocopied hypomorphic alleles in all our assays. Strains bearing fluorescently tagged septins exhibited defects in zygote cytokinesis, qualitatively phenocopying both hypomorphic and null alleles. These findings agreed with recent work with fission yeast, demonstrating the sensitivity of septin function to tagging. Interestingly, tagging septins did not perturb postembryonic development including animal mobility. This suggests that septins play distinct functions in the zygote versus later in development.

## Introduction

Septins are small GTP-binding proteins that form hetero-oligomeric complexes that assemble into cytoskeletal filaments and sheets. Septins contribute to cytokinesis, morphogenesis, and other cellular processes ^1,2^. Septins bind several components of the cortical cytoskeleton, including F-actin and NMMII. Septins also sense micron-scale membrane curvature ^3^. Septins have been implicated in many diseases, including cancer, infertility, congenital defects, and Alzheimer’s Disease, bipolar disorder, and other neurological disorders ^4,5^.

Despite their fundamental importance to basic cell functions, and implication in pathologies, septins are under studied and relatively poorly understood. These deficits stem, in part, from the large number of partially redundant septin genes in the genome of various model organisms including powerful fungal systems and all mammals. By contrast, the *C. elegans* genome has only two septin genes, *unc-59* and *unc-61*, which encode proteins that form a nonpolar heterotetramer ^6,7^. The *unc-61* locus encodes three isoforms: a, b, and c. *unc-61a* is transcribed from an alternative start site upstream of the shared start codon of the *unc-61b* and *unc-61c isoforms*, which differ only by a single stretch of four amino acids, a small difference that has not been functionally explored ^8^. As in some fungi and in other animals, in *C. elegans*, septins are enriched in the cytokinetic ring where they contribute to cytoskeletal scaffolding and organization. These roles manifest, in cells lacking septin function, in failed cortical rotation (reflecting torque release) and abnormally symmetric cytokinetic ring closure ^9-11^. That said, *C. elegans* septins are generally dispensable for cytokinesis except when contractility is compromised ^9^. Both *unc-59* and *unc-61* mutants generally have normal embryonic development; defects arise in post-embryonic development, affecting the morphogenesis of the vulva, male tail, gonad, and sensory neurons, and resulting in uncoordinated movement and reduced fecundity ^6^. While the two septins are believed to work together, there is evidence that each has unique roles in certain neurons and in germline development ^12,13^.

Fluorescent tagging is a powerful approach to study protein complex assembly and structure, localization, and function. However, it is not without issue. One popular probe for F-actin, Lifeact, partially inhibits actin assembly and constriction of the cytokinetic ring, and alters actin filament arrangements and dynamics, in fission yeast ^14^. Lifeact also impacts axon outgrowth ^15^. Similarly, C-terminal tagging of tubulin of alpha or beta tubulin with GFP impedes their interaction with microtubule-associated proteins; N-terminal GFP tagging of alpha or beta tubulin obstructs its incorporation into the microtubule lattice ^16-18^.

Green fluorescent protein originally isolated from the *Aequora Victoria* jellyfish can form low-affinity dimers under physiological conditions, particularly when fused to naturally oligomeric proteins or confined to membranes, leading to mistargeting and aggregation of fused constructs ^19^. Naturally occurring orange-and red fluorescent proteins isolated from other organisms are either dimeric or tetrameric. To overcome this issue, monomeric red fluorescent proteins have been engineered ^20^. One of these monomeric red proteins is a version of mScarlet that has been codon-optimized for expression in *C. elegans*: wrmScarlet ^21^. Monomeric red proteins are undesirably dim, with the brightest among them, mKate, being considerably dimmer than any naturally occurring red fluorescent protein. We used mKate2, which is more photostable and 3-fold brighter than mKate ^20^.

We found that fluorescent tagging disrupted the function of both septins. While septins are implicated in various roles within the cytoskeleton, we observed phenotypes specifically in zygotes and not during post-embryonic development. Results were comparable for tagged UNC-59 and UNC-61b/c, with various tags. Zygote cytoskeleton phenotypes included reduced cortical rotation during early anaphase, increased ring closure symmetry, and increased blebbing. The lack of septin phenotypes in larval and adult worms, and the normal localization of fluorescently tagged septins suggests that specific, limited septin roles, such as cytoskeletal scaffolding, are disrupted by the attachment of fluorescent proteins.

## Methods

### *C. elegans* strain and maintenance

*C. elegans* (see Table 1 for strain names and genotypes) worms were maintained on nematode growth medium (NGM) and OP50 bacterial food at 20°C.

### *C. elegans* DIC imaging conditions and analysis

Embryos were excised from adult worms. For each recording, cortical plane sections were separated into a series of TIFF image sequences using Fiji ^22,23^. Image sequences were analyzed using PIVlab, a particle image velocimetry software, a plugin for Matlab ^24^. From the resulting vector field of the velocities, vectors perpendicular to the cell long axis were separately averaged. Velocity is defined as the speed in the right-handed direction perpendicular to the anterior– posterior axis. Vectors were averaged within a region of interest measuring 13 microns in width and 35 microns in height placed within the center of the embryo. The program was calibrated to output velocities in microns per second.

### Image acquisition Cortical Rotation

Within a drop M9 buffer solution worm embryos were excised. Embryos were then transferred onto 2% agarose pads and a cover slip was placed on top. Images were acquired by DIC (Differential Image Contrast) microscopy using a Delta Vision Elite microscope (Cytiva) with an Olympus 60 × 1.40 NA oil immersion objective lens. For three minutes, two optical sections were acquired every two seconds: the midplane of the cell and the cortical plane proximal to the coverslip. The midplane was recorded to determine the timing of anaphase onset; the cortical plane was where granule movement was observed and quantified. Identical acquisition settings were used for all zygote imaging. To generate a true to life image orientation, images of *C. elegans* embryos were mirrored across the AP axis using FIJI tools ^11^.

### Particle Image Velocimetry analysis

Embryos were rotated so that the AP axis was oriented vertically with the anterior at the top of the image. Movies were separated into a sequence of image frames. Cortical granule displacement was measured between each pair of consecutive frames to determine rotational velocity. Average velocity was measured within a 35 × 13 micron region of interest centered along the cell’s AP axis (see Fig. 1B). Velocity was calculated by averaging the “U” or horizontal (circumferential) component of vectors within this region of interest during a 50 frame (100 second) time window that began 20 frames (40 seconds) after anaphase onset. Anaphase onset was determined by a sudden elongation of the spindle in the midplane region of the cell.

**Figure 1.**
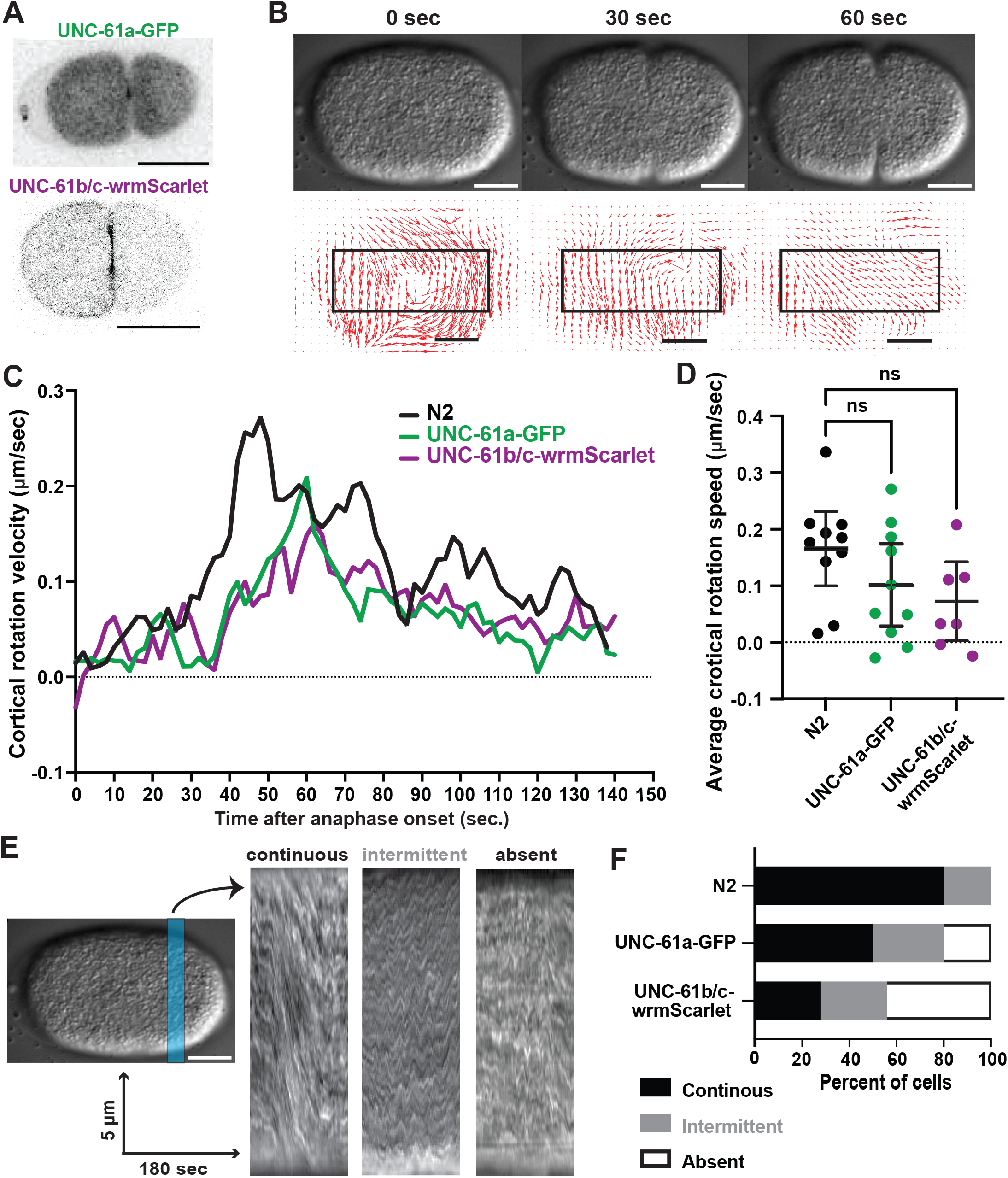
Fluorescent tagging of *C. elegans* septins causes dominant loss of septin function during anaphase cortical rotation in the zygote. **A)** Fluorescence images of UNC-61a-GFP and UNC-61b/c-wrmScarlet localization in zygotes late in cytokinesis. Scale bars = 20 µm. **B)** Top: DIC imaging at the coverslip-proximal plane of a control embryo showing cortical rotation; below: corresponding particle image velocimetry-generated velocity fields; arrow length represents local velocity. Box: 35 × 13 micron region for velocity measurements. Scale bars = 10 µm. **C)** Timepoint averaged circumferential (around the anterior–posterior axis) velocities during cortical rotation. **D)** Average circumferential cortical rotation velocities (bars: 95% confidence intervals). ANOVA statistical analysis results noted. **E)** DIC image of the coverslip-proximal cortical plane of a zygote; blue box: region for kymograph generation (left), and representative kymographs for each flow classification (right). **F)** Rotation phenotype frequencies. Throughout, N2 (n=10), UNC-61a-GFP (n=10), UNC-61b/c-wrmScarlet (n=7).

Cortical rotation classification was determined using average vector velocity measurements. Rotation was classified as continuous for cells in which there was at least one single period of at least 30 seconds of circumferential velocity greater than or equal to 0.05 microns per second (one quarter of the maximum speed). Intermittent was the score for cells with at least one period of 8-30 seconds with circumferential velocity > 0.05 microns/sec. A condition of none (no rotation) was assigned for cells that failed to meet either of these previously mentioned parameters.

### Kymograph Creation

Kymographs were made using FIJI’s multi-kymograph tool. A 10 pixel wide line was drawn perpendicular to the anterior-posterior axis such that the line originated from the top of the image, with anterior on the left (Fig. 1E).

### Ring Acquisition method

Within a drop M9 buffer solution, embryos were excised from gravid adult hermaphrodites. A square of VaLaP (1:1:1 mixture of Vaseline, lanolin, and paraffin) roughly the size of a cover slip was drawn to surround the M9 drop creating a rim slightly higher than the drop. A cover slip was then placed on top of the VaLaP rim allowing the cover slip and M9 drop to come into contact. The coverslip was then sealed by tracing its edges with VaLaP. The slide was then inverted allowing the embryo to settle onto the coverslip. Time lapse image series were acquired on a Nikon A1R microscope body with a Gallium arsenide phosphide photo-multiplier tube (GaAsP PMT) detector using a 60 × 1.27 NA water immersion objective lens and NIS-elements. Temporal resolution (TR) differed among conditions based on each tagged strain’s brightness and propensity for photobleaching. mNG-ANI-1: TR= 2 seconds. UNC-59-GFP: TR= 3 seconds A 5 μm area containing the cytokinetic furrow was selected and an end-on reconstruction was created using the reslice function in FIJI. The resulting image stack was average-intensity projected into two dimensions. The resulting tiff stack was then segmented using a custom Python script that fits a polygon to the cytokinetic ring based on mNeonGreen-ANI-1, or UNC-59-GFP fluorescence intensity. The resulting polygon was visually verified and divided into 72 equal angular segments and the coordinates for the center of the segment were recorded for each time point. The Fourier coefficients of the resulting coordinates were spatially smoothed by fitting a 2D harmonic series with 3 harmonics and temporally smoothed using a moving average over 5 time points. A smoothed inward velocity of individual segments was derived using a smooth differentiator function described previously ^25^.

### Ring Symmetry Measurement

Dividing *C. elegans* zygotes were imaged at the midplane using Nikon A1R scanning confocal 60 × 1.27 NA water immersion objective lens. Using FIJIs line tool the distance from the ring’s edge to the cell’s edge was measured when the cytokinetic furrow reached 50% closure. A ratio of these distances was then taken.

### Bleb analysis

A hypomorphic allele of *unc-59* served as a positive control, and, as the hypomorph strain bore a fluorescent plasma membrane marker, we used the parental membrane marker strain (OD95) for our negative control. Images of zygotes were acquired at the midplane for four minutes using DIC. Blebs were manually counted. In FIJI tools a segmented line was used to trace the perimeter of each bleb at its maximum expansion. The 2D area within this line was then quantified along with how many frames the bleb persisted.

### Brood Count

Fourth larval (L4) stage worms for each strain were singled onto individual plates. Worms were chosen from non starved growth plates and placed on individual cultures plates at 20 °C for 48 hours. After 24 hours, the then-adult worm was placed on a new individual culture plate and incubated at 20 °C for another 24 hours. After 48 hours, the adult worm was removed and all larvae and unhatched embryos were manually counted using a handheld tally counter tool and a Nikon SMZ800 dissection microscope.

### Embryonic Viability

Fourth larval (L4) stage worms from non-starved growth plates were placed singly on individual culture plates and incubated at 20 °C for 48 hours. After 24 hours, the then-adult worm was placed on a new individual culture plate and incubated at 20 °C for another 24 hours. Following each 24 hour period, the worm was removed and the number of eggs laid was counted. Following another 24 hour period, the number of viable worms was counted. A ratio was then taken comparing the number of viable worms to the number of eggs laid. This was done for both culture plates. After 48 hours, the adult worm was removed and all larvae and unhatched embryos were manually counted using a handheld tally counter tool and a Nikon SMZ800 dissection microscope.

### Phenotype Screen

100 µl of bleach solution (4 ml 5% bleach + 3.5 ml water + 2.5 ml 1M NaOH) was placed onto a fresh agar plate away from bacterial lawn. 30-50 adult worms were placed into the pool of bleach solution. Plates were placed agar side down so that bleach solution could evaporate leaving behind only the eggs. Worms were then placed into an incubator at 20 °C for 56 hours. Animals that had not reached adulthood (immature), and those exhibiting septin phenotypes (slight protruded vulva, strongly protruded vulva, and uncoordinated movement) were scored.

### Figures

Figures were generated using Microsoft Excel, Python, Matlab, PIVlab and Prism.

### Quantification and Statistical Analysis

Statistical significance was determined using a one-way ANOVA test for multiple comparison with the same control using corresponding Prism functions. A p value of less than

0.05 was considered significant. All error bars represent the mean with a 95% confidence interval. Sample Size (n) is indicated in the text or the figure legend.

### Software and algorithms

|  |  |  |
| --- | --- | --- |
| FIJI | Schindelin et al. | https://imagej.net/downloads |
| Matlab | Mathworks | https://www.mathworks.com/products/matlab.html |
| Python | Anaconda | https://www.anaconda.com/products/navigator |
| Microsoft Office | Microsoft |  |
| Prism | GraphPad | https://www.graphpad.com/features |
| PIVlab | Mathworks | https://www.mathworks.com/matlabcentral/fileexchange/27659-pivlab-particle-image-velocimetry-piv-tool-with-gui |
| Ring segmentation, Mode decomposition, data analysis and mathematical model | https://zenodo.org/records/11478957 | https://github.com/michaelewerner/Werner_et.al._submitted_2023 |

### C. elegans strains and culture

| strain | Genotype | Source or Reference |
| --- | --- | --- |
| N2 | Wild type |  |
| OD95 | ItIs37 [pAA64; pie-1::mCherry::HIS-58 + unc-119(+)] IV. ItIs38 [pAA1; pie-1::GFP::PH(PLC1delta1) + unc-119(+)]. Superficially wild type | <sup>26</sup> |
| MDX40 | mNeonGreen::ANI-1 x *nmy-2*::mKate2 CRISPR (LP229) | <sup>27</sup> |
| MDX113 | unc-59 [prt131 (*unc-59* KO)] I; *unc-119(ed3)* III (?) | <sup>28</sup> |
| MDX118 | mon22 *unc-61* KO CRISPR | <sup>8</sup> |
| JCC661 | unc-59(e1005) I.; unc-119(ed3) III?; ItIs38 [pAA1; pie-1/GFP::PH(PLCdelta1); unc-119 (+)] III.; unc-119 (+) III; ItIs37 [pAA64; pie-1/mCherry::his-58; unc-119 (+)] IV | <sup>11</sup> |
| NK2228 | *unc-59(qy88[unc-59::GFP-C1::3xflag::AID+loxP])* | <sup>29</sup> |
| COP1481 | COP1481 unc-59 (Knu463 - unc59::degron::mKate2) | <sup>30</sup> |
| COP2585 | knuSi920[pNU3401(AMAD02-unc-61p::UNC-61(a)::GFP::unc-61u+ Cbr-unc-119(+)] II;unc-119(ed3)III | <sup>8</sup> |
| COP2601 | knuSi934[pNU3400(AMAD01-unc-61p::UNC-61(B/C)::wrmScarlet::unc-61u, unc-119(+)]II; unc119(ed3)III | <sup>8</sup> |
| MDX141 | knuSi934 [pNU3400 (AMAD01- unc-61p::wrmScarlet::UNC-61(B/C)::unc-61u, unc-119(+))] II; unc-119(ed)III::unc-61(mon22 [unc-61 KO]) V | This work |

## Results

### Fluorescent tagging of septin isoforms slightly reduces cortical rotation

Fluorescent tagging is a powerful strategy for understanding protein dynamics and function. In the course of our recent work reporting novel functions of different isoforms of the *C. elegans* septins UNC-61a and UNC-61b/c, we generated strains expressing fluorescently tagged UNC-61a and UNC-61b/c under the control of their own promoters. These transgenes were inserted at an exogenous locus; the endogenous locus is wild type and intact. Here, we explored these isoforms’ localization and function within the zygote. Both isoforms were enriched in the cytokinetic ring (Fig. 1A), as is the endogenous protein and previously characterized fluorescently tagged septins ^6,9,29^. We next explored how exogenous fluorescently tagged septins, in the presence of wild type untagged septins, affect a septin-dependent process. Septins’ roles in scaffolding the cortical actomyosin cytoskeleton manifest in whole-cell cortical rotation (Fig. 1B) ^11^. Our analysis of cortical dynamics by particle image velocimetry revealed that rotation trended slower than wild type controls, but was not significantly different (Fig. 1C, D). As we have previously done, we classified cortical rotation phenotypes into continuous, intermittent, and absent, based on the duration of rotation faster than specified velocity thresholds (see Methods) ^11^. Cortical rotation was more likely to be intermittent or absent in cells expressing fluorescently tagged UNC-61a or UNC-61b/c than controls, suggesting that torsional forces did not accumulate in these cells to the same degree as in wild type (Fig. 1E, F). Thus, fluorescent tagging does not appreciably alter septin localization in *C. elegans*, as occurs in fission yeast ^31^, but causes mild dominant effects on the cortical cytoskeleton, mimicking loss of function alleles ^11^.

### Septin null strains exhibit cytoskeletal scaffolding defects

We concluded that fluorescent tagging perturbs septin function by comparing the tagging strains to hypomorphic alleles generated by random mutagenesis ^6,11^. Here, we sought to characterize full loss of septin function using null alleles of both *unc-59* and *unc-61* ^28^. Deletion of either *unc-59* or *unc-61* perturbed cortical rotation to a similar degree as the previously characterized septin hypomorphs (Fig. 2A, B) ^11^. Thus, as in the hypomorphs, in *unc-59* and *unc-61* knockouts, relief (and likely accumulation) of cytoskeletal torque in early anaphase is diminished or absent.

**Figure 2.**
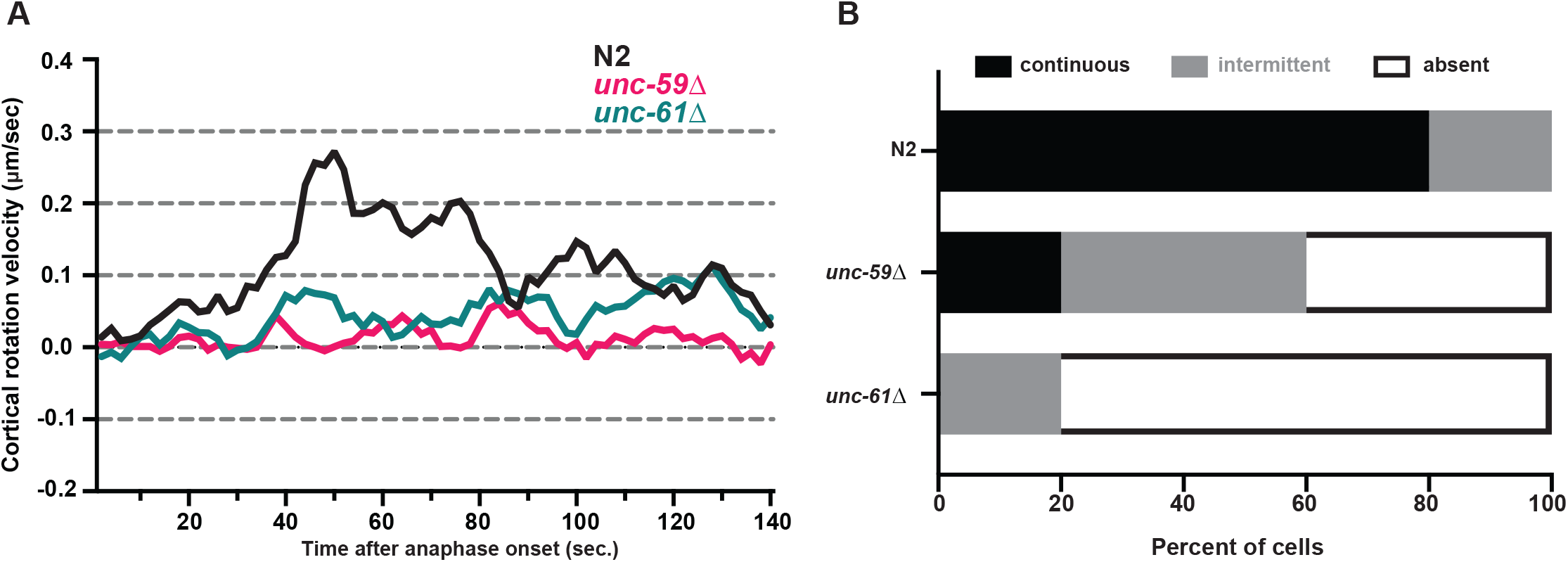
Septin deletion diminishes anaphase cortical rotation in the *C. elegans* zygote. **A)** Timepoint-averaged circumferential cortical rotation velocities. **B)** Rotation phenotype frequencies. N2 (n=10), *unc-59*Δ (n=10), *unc-61*Δ (n=10).

### Replacement of septins with fluorescently tagged transgenes perturbs cortical cytoskeletal network dynamics

Towards understanding what functions can be conferred by tagged septins in the absence of their endogenous counterparts, we next examined zygotes from strains wherein conventional GFP or mKate2 had been inserted at the endogenous *unc-59* locus ^29,30^. Endogenously-tagged UNC-59-GFP and UNC-59-mKate2 localized normally to the cytokinetic ring (Fig. 3A). To determine if the presence of fluorescent tags perturbed septins’ roles in the buildup of cytoskeletal torque during zygote anaphase, we again measured cortical rotation velocity using particle image velocimetry. In many zygotes expressing UNC-59-GFP or UNC-59-mKate2, cortical rotation had reduced velocity or was undetectable (Fig. 3B-D). Similarly, *unc-61(null)* zygotes with UNC-61b/c-wrmScarlet, cortical rotation was perturbed to a comparable degree as in the other tagged strains (Fig. 3C-E). We do note that the animals expressing UNC-61b/c-wrmScarlet were null for *unc-61a*, the isoform encoding the transmembrane domain, which may contribute to normal cortical dynamics in the zygote (Fig. 1B-E).

**Figure 3.**
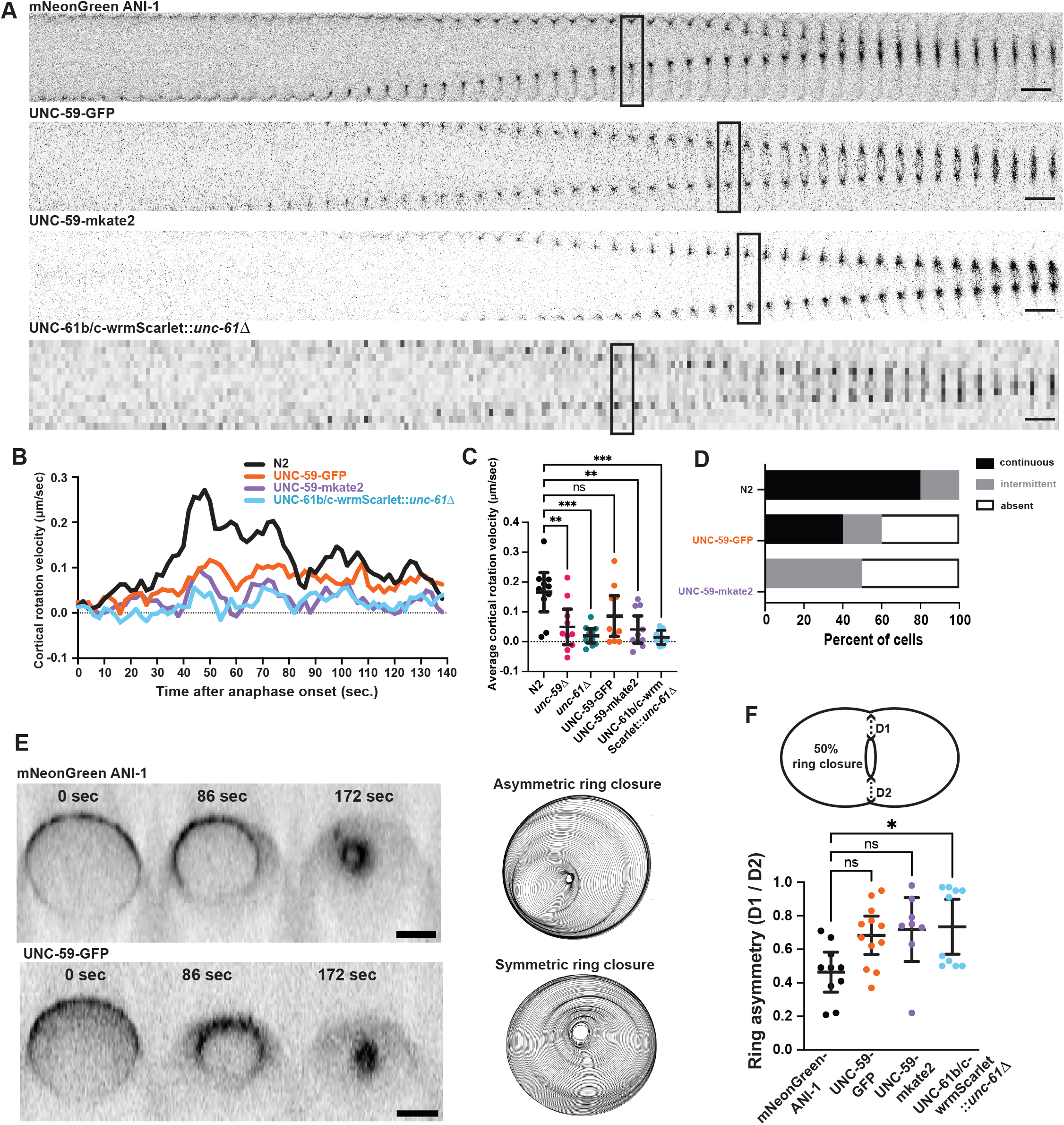
Tagging *C. elegans* septins with GFP or mKate2 perturbs cortical cytoskeletal dynamics in anaphase. **A)** Montage of equatorial region during cytokinetic furrowing for strains indicated. Boxed frames: ∼50% ring closure. **B)** Timepoint-averaged cortical rotation velocity. **C)** Average circumferential velocities during cortical rotation (bars: 95% confidence intervals). ANOVA p values * <0.005,**<0.01,*** <0.001. **D)** Frequency of rotation phenotype classifications. **E)** X, Z projections of 4D image series (right) and all-timepoint overlays (right) showing cytokinetic ring positioning within the division plane, over time, for fluorescent markers indicated. **F)** Diagram (top) and graph (bottom) of ring symmetry measurements (mean and 95% confident intervals); 1 = symmetric, 0 = unilateral. * *p* < 0.05. mNG-ANI-1 (n=10), UNC-59-GFP (n=12), UNC-59-mKate2 (n=8), UNC-61b/c-wrmScarlet::*unc-61*Δ (n=10). Scale bars = 10 µm.

Septins’ roles in crosslinking the actomyosin cytoskeleton and/or scaffolding it to the plasma membrane also manifest in ensuring the zygote cytokinetic ring’s asymmetric closure, a feature thought to optimize its energy efficiency ^10^. To assess whether fluorescently tagged septins confer this role, we compared the asymmetry of ring closure in our septin strains to that in a strain in which the fluorescent tag does not perturb ring asymmetry (ANI-1-anillin-GFP) ^27^. Cells in which UNC-59 or UNC-61b/c were replaced with UNC-59-GFP or UNC-59-mKate2, or UNC-61b/c-wrmScarlet, exhibited more symmetric ring closure than controls (Fig. 3A, E, F). However, decreased asymmetry was only statistically significantly different for UNC-61b/c-wrmScarlet, which had a bimodal distribution of asymmetry and which lacks UNC-61a (Fig. 3A, E, F).

In sum, while fluorescent tagging of *C. elegans* septins does not perturb septin localization as occurs in fission yeast ^31^, it does perturb septins’ roles in cytoskeletal scaffolding that manifest in zygote cortical dynamics.

### Replacement of septins with fluorescently tagged transgenes causes blebbing

Anchoring the actomyosin cytoskeleton to the plasma membrane, septins help prevent blebbing, which occur when intracellular hydrostatic pressure overcomes membrane-cytoskeleton coupling ^32^. We explored whether the deleterious effects of fluorescent tagging on septin function perturb cortical stability, causing blebs. Blebs were seldom observed in our negative control cells (Fig. 4). Embryos bearing a hypomorphic allele of *unc-59* exhibited frequent blebbing both in the cell equator flanking the cytokinetic furrow (Fig. 4A) and at the anterior pole (Fig. 4B). To test the effects of tagging UNC-59, we examined strains in which either GFP or mKate2 had been introduced at the endogenous *unc-59* locus. Both strains exhibited much more blebbing than our negative control (Fig. 4). Interestingly, blebbing severity varied depending on which tag was used. Cells with UNC-59-mKate2 blebbed more than cells with UNC-59-GFP (Fig. 4). We also assessed the effect of fluorescently tagging UNC-61b/c on membrane-cytoskeleton coupling by measuring blebbing in our strain in which tagged UNC-61b/c is the only source of this septin (but in which UNC-61a is absent). Cells with wrmScarlet-tagged UNC-61b/c had fewer blebs than our unc-59 mutant control and cells with tagged UNC-59 (Fig. 4C-F). Together, these results further support the idea that tagging septins perturbs their function, though the severity of perturbation varies among tagged-septin conditions.

**Figure 4.**
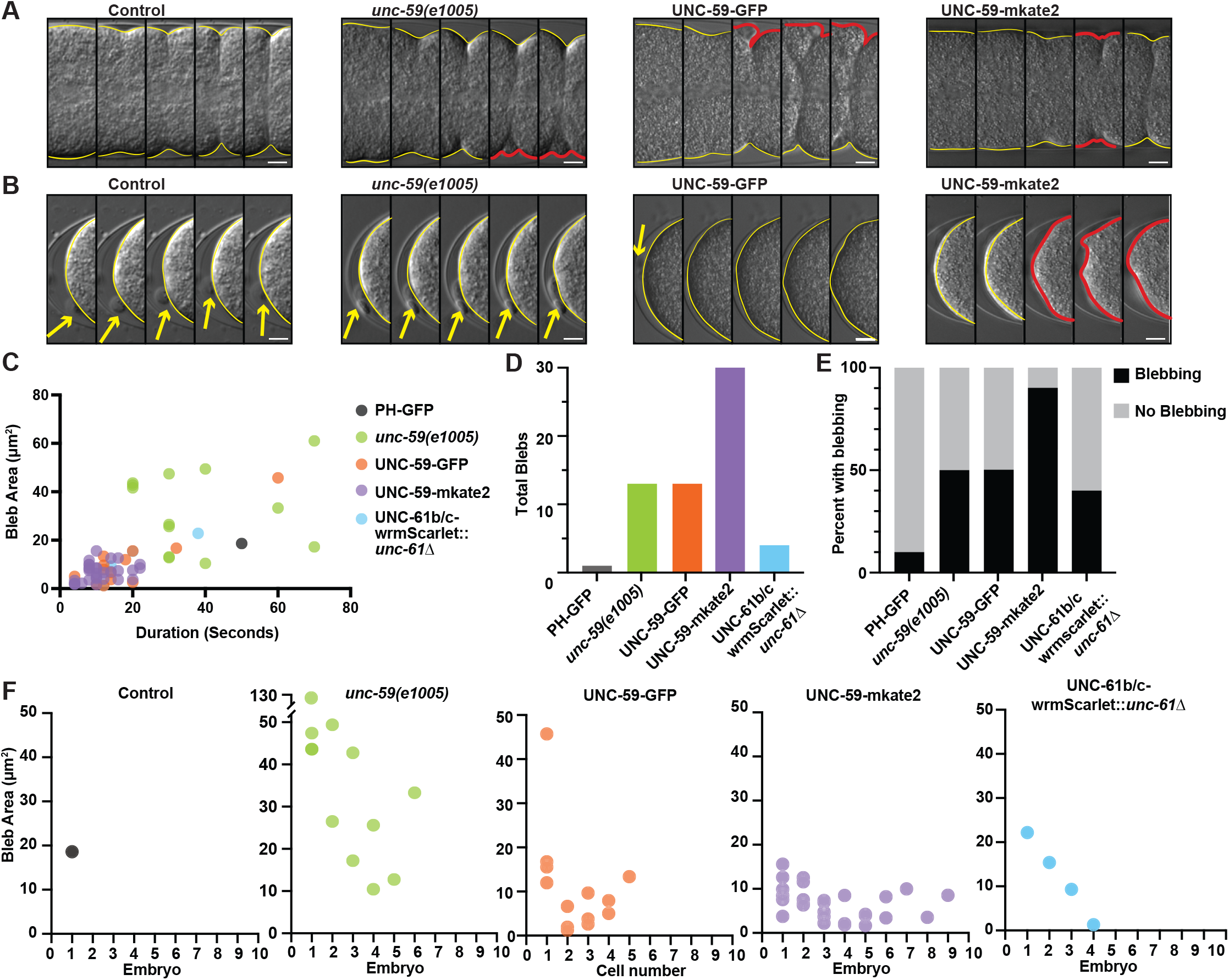
Tagging *C. elegans* septins with GFP or mKate2 increases blebbing. **A)** Montage of the cytokinetic furrow region in a negative control and the other genotypes indicated. **B)** Montage of zygote anterior during anaphase. (**A & B** yellow outlines: quiescent cortex or normal furrow; red outlines: blebs; yellow arrows: polar body; temporal resolution 40 or 50 seconds; scale bars =10 µm. **C)** Bleb area plotted as a function of duration. **D)** Total number of blebs observed in all embryos. **E)** Percent of embryos that exhibited blebbing. **F)** Bleb quantity and area experienced by each individual embryo. (**C-F** n = 10 embryos per strain strain).

### Fluorescently tagged septins can support development

Fluorescent tagging of septins phenocopied septin mutants in our assays of actomyosin cortical dynamics and membrane anchoring in the early embryo. However, anecdotally, our strains bearing tagged septins appeared relatively normal throughout routine maintenance, when observed via stereoscope. To test whether the same conditions that perturbed septin function in early embryogenesis affected other events, we scored several post-embryonic phenotypes in which septins have been implicated ^6^.

First, we measured the number of progeny produced by animals of various genotypes, in the first two days of adulthood. The fecundity of animals in which all UNC-59 was tagged with GFP was statistically indistinguishable from controls. By contrast, fluorescent tagging of UNC-59 with mKate2 did reduce fecundity, though not to the extent of *unc-59* deletion (Fig. 5A). Similarly, animals with UNC-61b/c-wrmScarlet as their only source of UNC-61 had lowered fecundity, but produced almost an order of magnitude more progeny than *unc-61(null)* animals, on average. We cannot eliminate the fertility defects exhibited by the latter strain stem from the absence of UNC-61a. In sum, these results suggest that tagging septins with fluorescent proteins can mildly perturb germline function.

**Figure 5.**
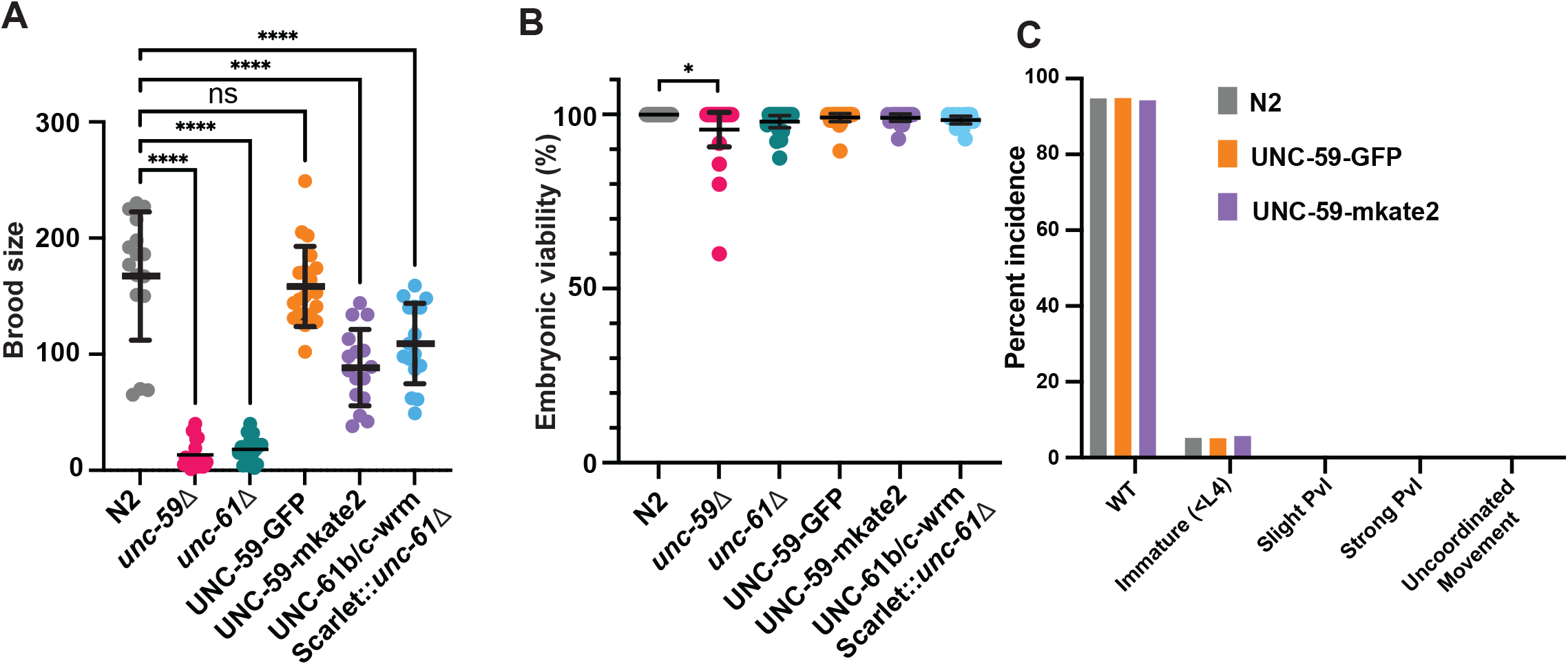
Tagging *C. elegans* septins with GFP or mKate2 reduces fecundity but not on embryonic viability or post-embryonic development. **A)** Number of eggs laid by individual animals within 48 hours. N2 (n=19), *unc-59*Δ (n=20), *unc-61*Δ (n=19), UNC-59-GFP (n=19), UNC-59-mKate2 (n=16), UNC-61b/c-wrmScarlet::*unc-61*Δ (n=16). **B)** Ratio of viable embryos versus total eggs laid in 48 hours. N2 (n=20), *unc-59*Δ (n=20), *unc-61*Δ (n=19), UNC-59-GFP (n=19), UNC-59-mKate2 (n=16), UNC-61b/c-wrmScarlet::*unc-61*Δ (n=16). **C)** Percent incidence of septin phenotypes for animals aged 24 hours beyond L4. N2 (n=385), UNC-59-GFP (n=412), UNC-59-mKate2 (n=421).

Septins are required for several cell biological events to be quantitatively normal ^6,11^. Interestingly, these roles appear to be dispensable for embryonic viability, since when septin function was perturbed via null mutation of *unc-59*, embryonic viability was only slightly lower than controls. *unc-61(null)* animals and those in which UNC-59 or UNC-61b/c was fluorescently tagged had embryonic viability statistically indistinguishable from controls (Fig. 5B). Thus, the defects we observed in the zygote did not translate into significant fitness disadvantages for the embryo for the animal.

To assess the effect of tagging septins on post-embryonic development, we synchronized animals at the fourth larval stage and assessed their morphology, mobility, and maturation after 24 hours. Loss of septin function is well known to cause uncoordinated movement (hence their gene names) and abnormalities of epithelial morphogenesis leading of protruded vulva, attenuated egg laying, and extrusion of the germline ^6^. We observed that by contrast, animals with endogenously fluorescently tagged UNC-59 exhibited none of these post-embryonic development phenotypes. Like our wild type controls, ∼93% of animals in tagged-septin strains had reached adulthood (Fig. 5C). (The requirement of UNC-61a for vulval morphogenesis precluded using our strain expressing UNC-61b/c-wrmScarlet on the background of *unc-61a(null)* for this same assay ^8^. Together, these results demonstrate that tagging septins preserves their function in a manner compatible with embryonic and post-embryonic development and fertility.

## Discussion

We examined the effects of tagging *C. elegans* septins with fluorescent proteins. We found that fluorescently tagged UNC-61a or UNC-61b/c, in the presence of wild type UNC-61, perturbed the dynamics of the cortical actomyosin cytoskeleton in the zygote. Furthermore, replacement of UNC-59 or UNC-61b/c with a fluorescently tagged version by endogenous tagging or via insertion at an exogenous site all perturbed septin function in the zygote, reducing cortical rotational velocity and increased ring closure symmetry and blebbing. By contrast, while septin null alleles resulted in severe sterility and rare embryonic lethality, post-embryonic development and tissue function was generally normal in strains bearing fluorescently tagged septins.

Since the version of GFP we used can form dimers ^19^, we had predicted that strains expressing UNC-59-GFP would exhibit more severe loss of function than those with UNC-59-mKate2, which was engineered to be monomeric, from the same locus ^20^. However, we observed the opposite. It is possible that artifactual multimerization of septins does not perturb septin function since septins themselves polymerize, or that GFP does not robustly form dimers in *C. elegans*. Our findings that monomeric fluorescent tags perturb septin function agree with a recent report about fission yeast septins. In fission yeast, monomeric red fluorescent protein tags more severely perturbed localization and function than did green fluorescent proteins retaining or relieved of the propensity to form dimers, or even small tags including HA ^31^. Future work will directly test how fluorophore multimerization affects animals septin, using *C. elegans*, by genetically altering GFP to eliminate dimerization ^33^.

Our findings presented here suggest UNC-61a is active in the zygote, suggesting that well characterized events including blebbing and cytokinetic furrowing can serve as contexts in which to explore the contributions of septins’ transmembrane domains, which occur widely throughout phylogeny but not in any other standard model organisms ^8^. Future work will be aimed at studying UNC-61a localization and dynamics throughout the zygote cell cycle, and testing whether these factors and cortical dynamics depend on the transmembrane domain.

We conclude that fluorescent tagging disrupts septins’ cytoskeletal scaffolding roles, as it slightly decreased cortical rotation velocity and ring asymmetry, and increased blebbing. Interestingly, septin tagging did not cause postembryonic phenotypes characteristic of loss of septin function, such as delayed development, vulval protrusion, or uncoordinated movement ^6^. Septins are required for post-embryonic neuroblast mitosis and axon guidance and migration ^12^, which rely on cytoskeletal scaffolding. Our findings thus contrast the prediction that the zygote and post-embryonic cells would have the same sensitivity to tagging septin. Instead, our findings suggest that septins’ post-embryonic roles are distinct from their contributions to membrane-cytoskeletal linkage and cytoskeletal crosslinking in the zygote. Other activities could include scaffolding signaling partners and conferring membrane subdomains, as septins do in other cell types ^34-36^. Future studies will explore the cell biological bases of septins’ roles in somatic cells including neuroblasts.

## Acknowledgements

The authors thank all members of the Maddox labs, especially Michael Werner, for assistance, advice, and critical reading of this manuscript. Arittri Mallick and Stephan Grill provided support with particle image velocimetry. Ana Filipa Sobral and Ana Carvalho provided the *unc-59* deletion strain. This work was supported by NIGMS R35GM144238 and NSF 2153790 to ASM.

## Movie Legends

**Movie 1. Septin deletion diminishes anaphase cortical rotation in the *C. elegans* zygote**. Time lapse of cells exhibiting **A)** continuous anaphase cortical rotation in N2 embryo, **B)** absent anaphase cortical rotation in *unc-59*Δ embryo, **C)** absent anaphase cortical rotation in *unc-61*Δ embryo.

**Movie 2. Tagging *C. elegans* septins with GFP or mKate2 perturbs cortical cytoskeletal dynamics in anaphase**.

Time lapse of cells exhibiting **A)** continuous anaphase cortical rotation in N2 embryo, **B)** absent anaphase cortical rotation in UNC-59-GFP embryo and **C)** absent anaphase cortical rotation in UNC-59-mKate2.

**Movie 3. Tagging *C. elegans* septins with GFP or mKate2 increases blebbing**

Time lapse of cell exhibiting **A)** absence of blebbing during cytokinesis in GFP-PH embryo, **B)** increased blebbing during cytokinesis in UNC-59-mKate2 embryo, **C)** increased blebbing during cytokinesis in *unc-59(e1005)* embryo, **D)** increased blebbing during cytokinesis in UNC-59::GFP embryo, **E)** mild blebbing in UNC-61b/c-wrmScarlet::*unc-61*Δ embryo.

## References Cited

1. Mostowy, S., and Cossart, P. (2012). Septins: the fourth component of the cytoskeleton. Nat Rev Mol Cell Biol 13, 183–194. 10.1038/nrm3284.

2. Bridges, A.A., and Gladfelter, A.S. (2015). Septin Form and Function at the Cell Cortex. J Biol Chem 290, 17173–17180. 10.1074/jbc.R114.634444.

3. Bridges, A.A., Jentzsch, M.S., Oakes, P.W., Occhipinti, P., and Gladfelter, A.S. (2016). Micron-scale plasma membrane curvature is recognized by the septin cytoskeleton. J Cell Biol 213, 23–32. 10.1083/jcb.201512029.

4. Pissuti Damalio, J.C., Garcia, W., Alves Macedo, J.N., de Almeida Marques, I., Andreu, J.M., Giraldo, R., Garratt, R.C., and Ulian Araujo, A.P. (2012). Self assembly of human septin 2 into amyloid filaments. Biochimie 94, 628–636. 10.1016/j.biochi.2011.09.014.

5. Das, A., and Kunwar, A. (2025). Septins: Structural Insights, Functional Dynamics, and Implications in Health and Disease. J Cell Biochem 126, e30660. 10.1002/jcb.30660.

6. Nguyen, T.Q., Sawa, H., Okano, H., and White, J.G. (2000). The C. elegans septin genes, unc-59 and unc-61, are required for normal postembryonic cytokineses and morphogenesis but have no essential function in embryogenesis. J Cell Sci 113 Pt 21, 3825–3837. 10.1242/jcs.113.21.3825.

7. John, C.M., Hite, R.K., Weirich, C.S., Fitzgerald, D.J., Jawhari, H., Faty, M., Schlapfer, D., Kroschewski, R., Winkler, F.K., Walz, T., et al. (2007). The Caenorhabditis elegans septin complex is nonpolar. EMBO J 26, 3296–3307. 10.1038/sj.emboj.7601775.

8. Perry, J.A., Werner, M.E., Omi, S., Heck, B.W., Maddox, P.S., Mavrakis, M., and Maddox, A.S. (2025). Animal septins contain functional transmembrane domains. Curr Biol 35, 1910–1917 e1915. 10.1016/j.cub.2025.03.004.

9. Maddox, A.S., Lewellyn, L., Desai, A., and Oegema, K. (2007). Anillin and the septins promote asymmetric ingression of the cytokinetic furrow. Dev Cell 12, 827–835. 10.1016/j.devcel.2007.02.018.

10. Dorn, J.F., Zhang, L., Phi, T.-T., Lacroix, B., Maddox, P.S., Liu, J., and Maddox, A.S. (2016). A theoretical model of cytokinesis implicates feedback between membrane curvature and cytoskeletal organization in asymmetric cytokinetic furrowing. Molecular Biology of the Cell 27, 1286–1299. 10.1091/mbc.E15-06-0374.

11. Zaatri, A., Perry, J.A., and Maddox, A.S. (2021). Septins and a formin have distinct functions in anaphase chiral cortical rotation in the Caenorhabditis elegans zygote. Mol Biol Cell 32, 1283–1292. 10.1091/mbc.E20-09-0576.

12. Finger, F.P., Kopish, K.R., and White, J.G. (2003). A role for septins in cellular and axonal migration in C. elegans. Dev Biol 261, 220–234. 10.1016/s0012-1606(03)00296-3.

13. Perry, J.A., Werner, M.E., Rivenbark, L., and Maddox, A.S. (2023). Caenorhabditis elegans septins contribute to the development and structure of the oogenic germline. Cytoskeleton (Hoboken) 80, 215–227. 10.1002/cm.21763.

14. Courtemanche, N., Pollard, T.D., and Chen, Q. (2016). Avoiding artefacts when counting polymerized actin in live cells with LifeAct fused to fluorescent proteins. Nat Cell Biol 18, 676–683. 10.1038/ncb3351.

15. Patel, S., Fok, S.Y.Y., Stefen, H., Tomanic, T., Paric, E., Herold, R., Brettle, M., Djordjevic, A., and Fath, T. (2017). Functional characterisation of filamentous actin probe expression in neuronal cells. PLoS One 12, e0187979. 10.1371/journal.pone.0187979.

16. Xu, K., Li, Z., Mao, L., Guo, Z., Chen, Z., Chai, Y., Xie, C., Yang, X., Na, J., Li, W., and Ou, G. (2024). AlphaFold2-guided engineering of split-GFP technology enables labeling of endogenous tubulins across species while preserving function. PLoS Biol 22, e3002615. 10.1371/journal.pbio.3002615.

17. Nishida, K., Tsuchiya, K., Obinata, H., Onodera, S., Honda, Y., Lai, Y.C., Haruta, N., and Sugimoto, A. (2021). Expression Patterns and Levels of All Tubulin Isotypes Analyzed in GFP Knock-In C. elegans Strains. Cell Struct Funct 46, 51–64. 10.1247/csf.21022.

18. Honda, Y., Tsuchiya, K., Sumiyoshi, E., Haruta, N., and Sugimoto, A. (2017). Tubulin isotype substitution revealed that isotype combination modulates microtubule dynamics in C. elegans embryos. J Cell Sci 130, 1652–1661. 10.1242/jcs.200923.

19. Wang, X., Song, K., Li, Y., Tang, L., and Deng, X. (2019). Single-Molecule Imaging and Computational Microscopy Approaches Clarify the Mechanism of the Dimerization and Membrane Interactions of Green Fluorescent Protein. Int J Mol Sci 20. 10.3390/ijms20061410.

20. Shcherbo, D., Murphy, C.S., Ermakova, G.V., Solovieva, E.A., Chepurnykh, T.V., Shcheglov, A.S., Verkhusha, V.V., Pletnev, V.Z., Hazelwood, K.L., Roche, P.M., et al. (2009). Far-red fluorescent tags for protein imaging in living tissues. Biochem J 418, 567–574. 10.1042/BJ20081949.

21. Cao, W.X., Merritt, D.M., Pe, K., Cesar, M., and Hobert, O. (2024). Comparative analysis of new mScarlet-based red fluorescent tags in Caenorhabditis elegans. Genetics 228. 10.1093/genetics/iyae126.

22. Schindelin, J., Arganda-Carreras, I., Frise, E., Kaynig, V., Longair, M., Pietzsch, T., Preibisch, S., Rueden, C., Saalfeld, S., Schmid, B., et al. (2012). Fiji: an open-source platform for biological-image analysis. Nat Methods 9, 676–682. 10.1038/nmeth.2019.

23. Rueden, C.T., Schindelin, J., Hiner, M.C., DeZonia, B.E., Walter, A.E., Arena, E.T., and Eliceiri, K.W. (2017). ImageJ2: ImageJ for the next generation of scientific image data. BMC Bioinformatics 18, 529. 10.1186/s12859-017-1934-z.

24. Thielicke, W., and Sonntag, R. (2021). Particle Image Velocimetry for MATLAB: Accuracy and enhanced algorithms in PIVlab. J Open Res Softw 9.

25. Werner, M.E., Ray, D.D., Breen, C., Staddon, M.F., Jug, F., Banerjee, S., and Maddox, A.S. (2024). Mechanical and biochemical feedback combine to generate complex contractile oscillations in cytokinesis. Curr Biol 34, 3201–3214 e3205. 10.1016/j.cub.2024.06.037.

26. Audhya, A., Hyndman, F., McLeod, I.X., Maddox, A.S., Yates, J.R., 3rd, Desai, A., and Oegema, K. (2005). A complex containing the Sm protein CAR-1 and the RNA helicase CGH-1 is required for embryonic cytokinesis in Caenorhabditis elegans. J Cell Biol 171, 267–279. 10.1083/jcb.200506124.

27. Rehain-Bell, K., Love, A., Werner, M.E., MacLeod, I., Yates, J.R., 3rd, and Maddox, A.S. (2017). A Sterile 20 Family Kinase and Its Co-factor CCM-3 Regulate Contractile Ring Proteins on Germline Intercellular Bridges. Curr Biol 27, 860–867. 10.1016/j.cub.2017.01.058.

28. Sobral, A.F. (2022). Functionalities and interplay of actin-filament crosslinkers in vivo. PhD (University of Porto).

29. Chen, D., Hastie, E., and Sherwood, D. (2019). Endogenous expression of UNC-59/Septin in C. elegans. MicroPubl Biol 2019. 10.17912/micropub.biology.000200.

30. Priti, A., Ong, H.T., Toyama, Y., Padmanabhan, A., Dasgupta, S., Krajnc, M., and Zaidel-Bar, R. (2018). Syncytial germline architecture is actively maintained by contraction of an internal actomyosin corset. Nat Commun 9, 4694. 10.1038/s41467-018-07149-2.

31. Gregory, J.R., Ricottilli, N.J., and Wu, J.Q. (2025). Localizations of the septin Spn4 tagged with GFP and mEGFP in fission yeast. MicroPubl Biol 2025. 10.17912/micropub.biology.001930.

32. Gilden, J.K., Peck, S., Chen, Y.C., and Krummel, M.F. (2012). The septin cytoskeleton facilitates membrane retraction during motility and blebbing. J Cell Biol 196, 103–114. 10.1083/jcb.201105127.

33. von Stetten, D., Noirclerc-Savoye, M., Goedhart, J., Gadella, T.W., Jr., and Royant, A. (2012). Structure of a fluorescent protein from Aequorea victoria bearing the obligate-monomer mutation A206K. Acta Crystallogr Sect F Struct Biol Cryst Commun 68, 878–882. 10.1107/S1744309112028667.

34. Estey, M.P., Di Ciano-Oliveira, C., Froese, C.D., Bejide, M.T., and Trimble, W.S. (2010). Distinct roles of septins in cytokinesis: SEPT9 mediates midbody abscission. J Cell Biol 191, 741–749. 10.1083/jcb.201006031.

35. Benoit, B., Pous, C., and Baillet, A. (2023). Septins as membrane influencers: direct play or in association with other cytoskeleton partners. Front Cell Dev Biol 11, 1112319. 10.3389/fcell.2023.1112319.

36. Radler, M.R., and Spiliotis, E.T. (2022). Right place, right time - Spatial guidance of neuronal morphogenesis by septin GTPases. Curr Opin Neurobiol 75, 102557. 10.1016/j.conb.2022.102557.

